# Role of pre and post interventions on cervical cancer knowledge levels among women students at the University of Gondar, Gondar, Ethiopia

**DOI:** 10.1101/492611

**Authors:** Meera Indracanti, Nega Berhane, Tigist Minyamer

**Affiliations:** Assistant professor, Institute of Biotechnology, University of Gondar, Gondar, Ethiopia. Email i.d.; Professor, Institute of Biotechnology, University of Gondar, Gondar, Ethiopia. Email i.d.

**Author notes:** Corresponding authors /.

## Abstract

**Background:** Cervical cancer is the second most common cancer in women aged 44 years and above in Ethiopia. Lack of awareness about the disease, lack of screening programs and inadequacy of vaccination in most regions of Ethiopia alarmingly increasing Human papillomavirus (HPV) infections and incidence of the disease. Educational intervention is a fast and effective primary preventive step to reduce the cervical cancer burden.

**Objective:** The present study was carried out to understand the impact of knowledge-based intervention and factors influencing the knowledge levels on young women attending college education at University of Gondar (UoG), Gondar.

**Method:** A cross-sectional comparative study was conducted and data was collected using a standardized self-administered questionnaire in both English and Amharic (Ethiopian main official language) and analysed using the Statistical Package for the Social Sciences software (SPSS ver.23, IBM).

**Results:** There was an increase in overall awareness about cervical cancer (symptoms, risk factors, screening methods, and vaccination) in all post intervened students compared to baseline knowledge levels (before education intervention) statistically at p<0.001 significance level. The mean age of the study participants was 20.86 years (SD, 1.86). Out of total 283 women student participants, overall baseline awareness about cervical cancer symptoms (81.6%, p<0.002), risk factors (94.8%, p<0.001), HPV (60.6%, p<0.001), screening (84.3%, p<0.001) and HPV vaccines (42.1%, p<0.001) was more in 4^th^ year and above over other respondents. After the intervention, knowledge levels increased in students 3^rd^ and above years over 1^st^ and 2^nd^-year students irrespective of the branch they belong. Initial awareness on various broad issues was 8.77 and after education intervention, it was 30.39 with mean overall knowledge increase of 21.62. However, baseline awareness was better on risk factors and poor on vaccination. After education intervention, an increase of 246% in overall knowledge about cervical cancer including symptoms, risk factors, HPV, screening and vaccination. Age, year of study, branch of study and family income were the explanatory variables significant on overall baseline knowledge levels and after education intervention, year of study was the only independent variable significant for the overall increase in knowledge levels.

**Conclusion:** The present study suggests that educational intervention as the primary preventive method is effective and young trained women volunteers belong both rural and urban areas will be important stakeholder to increase positive attitude to reduce the cervical cancer burden in Ethiopia.

## Introduction

According to GLOBOCAN 2018 [1], most of the African countries have no official registry to cover the cancer statistics and it reflects unseen burden including cervical cancer. Cervical cancer is a fourth leading cause for cancer death is the most common cancer in Sub-Saharan Africa, second leading health problem in Northern Africa including Ethiopia among women 44 years and above [2–10]. In developing countries, high-risk HPV infections cause cervical cancer and other serious public health problems [11] due to bare minimal resources to cope with the situation [12–14].

Women are at risk of HPV infections in some point in their life [4, 15]. A variety of clinic-epidemiological risk factors such as early age of marriage, multiple sexual partners, multiple pregnancies, poor genital hygiene and smoking and so are often associated with the development of cervical cancer [4, 11, 16, 17]. Most of the women in developing and under-developed countries do not have access to Pap (Papanicolaou) smear screening [12, 18] for early detection of HPV infections. Low or absence of any nationwide cervical screening program [19], very few women receive screening [20] and cancer of cervix remains a major public health problem for Ethiopia [21, 22]. According to Tsegaye et al., 2018 [2], only 0.6% of women in Ethiopia, aged 18-69 years includes, 1.6% from urban and 0.4% from rural screened every three years. In Ethiopia, every year around 7095 women are diagnosed with cervical cancer and 4732 dies from this disease [23].

Several factors like education, economic status, health facilities influence early detection and treatment of cervical precancerous lesions [5, 15, 24–26] and reduce cervical cancer morbidity and mortality [27]. The absence of screening facilities coupled with poor literacy and low level of awareness, less attention to women health further aggravate the cervical cancer burden [5, 28–30]. Ethiopia has a low level of awareness about cervical cancer and HPV infections [27]. Various studies [31–33] have been undertaken to assess women’s awareness and knowledge level about cervical cancer. Cervical cancer awareness studies are few in Ethiopia and mostly confined to hospitals [4, 12, 34]. A recent study on women in the Amhara region has a low level of awareness [5] and factors influence the levels of knowledge not well known [4].

The success and benefit of control and prevention of cervical cancer largely depend to a great extent on the level of awareness and knowledge about different aspects of the disease and the vaccine [35, 36] and current focus on risk factors will be beneficial [37] and effective. It is therefore beneficial to understand the baseline knowledge levels of young women, awareness, and attitude towards cervical cancer and factors influencing their knowledge levels before and after education intervention towards effective primary preventive measure for control of cervical cancer burden in Ethiopia. Recent years, few studies were carried out to understand the baseline knowledge levels at the community level and as well at the university level in some parts of Ethiopia [2] and no study carried out to measure the knowledge levels and influence of socio-demographic factors before and after the educational intervention. So, the aim of this study was to explore cervical cancer knowledge levels of the students from two campuses of University of Gondar (UoG) and influence of any socio-demographic parameters on overall knowledge levels of study participants before and after the educational intervention.

## Materials and methods

### Study area and subjects

A cross-sectional pre-test/post-test comparative study was conducted to understand the socio-demographic factors (Independent variables (IVs)) influence on knowledge levels of women students of biological and non-biological sciences from Tewodros and Marakhi campuses of UoG. These two campuses have colleges for Computational & Natural Sciences and Management & Economics. The study included written informed consent and data collection tool was approved by the Department Research Committee, Institute of Biotechnology, UoG. Most of the students were from different regions of Amhara, Addis Ababa, Oromia and Southern Nations, mostly from rural areas belong to less educated families with less access to print and visual media.

### Sample size and questionnaire

#### Sample size

In UOG, the number of female students enrols to different programs is usually a low and average ratio of one female student to five male students. Any women aged 17 to 30 years enrolled in university graduate or postgraduate programs were invited to participate in the study. The study was conducted in a total of 283 undergraduate and postgraduate female students aged between 17–30 years. Based on the pilot study, the sample size was calculated using a formula for finite population [38]. The assumption was 50% of the university students had sufficient knowledge of cervical cancer, a sample of 283 students was selected by stratified random sampling techniques with 95% confidence and 5% reliability. Respondents were enrolled using a multistage sampling technique. Enrolled female students with eligible age volunteered to participate and signed written consent form were included in the study.

#### Questionnaire development

The questionnaire was designed and developed based on study objectives, literature review, and pilot study. An initial pilot study was carried out from May-June 2017 at the University of Gondar, Tewodros and Marakhi campuses, to test the data collection tool in English includes seven sections with 78 questions. During September-February’ 2018, the study was carried out using modified data collection tool consists of seven sections include 56 items both open- and close-ended questions in English and Amharic languages as most students preferred to use the questionnaire in Amharic.

The six-part questionnaire included socio-demographic characteristics and questions regarding the knowledge about different aspects of cervical cancer like: (1) Demographic characteristics, such as age, sex, religion, biological or non-biological sciences as study background, place of residence, father’s and mother’s educational qualifications and occupation, family size, family income of the students. (2) Awareness and knowledge of cervical cancer symptoms, (3) Knowledge of risk factors, (4) Knowledge of HPV, (5) Knowledge of cervical cancer screening, (6) Awareness and knowledge about HPV vaccine and awareness and perception towards screening, concern/acceptability of vaccination, health-seeking behaviour and preferences of venue for screening and vaccination.

Categorical data on various socio-demographic factors, continuous data on family income and age were collected. The purpose and importance of the study were explained to the participants prior to obtaining written informed consent and the confidentiality of their identities was ensured. The questionnaire was administered to the female students and the data from the questionnaire was processed anonymously by assigning random codes. Confidentiality of the information was maintained throughout by excluding names or I.D. Nos. in the questionnaire during data collection. Students were categorized into groups based on different factors, in order to examine which socio-demographic factors were strongly associated with the knowledge, awareness, and attitudes towards cervical cancer, HPV and vaccination. According to age, students were divided into two categories: young females aged 17 to 20 years and adult females aged 21 years & above. The education level of the students was classified into four groups: (i) first year, (ii) the second year, (iii) the third year, and (iv) fourth year & above. The household income per month was an open-ended question and based on the response it was classified into three categories as follows (i) <2000 birr (ii) >=2000-5000 birr (iii) >5000 birr and above. Knowledge levels of respondents regarding symptoms, risk factors, HPV and its relationship with cervical cancer, prevention methods like screening and vaccination was measured using a 42 item instrument. A score of 1 was allocated for a good/correct answer and 0 for a wrong answer or “Do not know”. The maximum possible score was 42. Mean score used to estimate the cumulative mean score of knowledge levels of cervical cancer. The total score was divided into, those scored above 31 or more were categorized as having very good (“sufficient”) knowledge; the others were categorized “good NK” with 23 to 31, fair with score 13-22 and poor NK was 1-12 and zero score categorized as “no” knowledge. Source of information, awareness and perception, concern and acceptability, health-seeking behaviour and choice of venue for screening and vaccination were measured before and after educational intervention and descriptive statistics were used to measure the change in response.

### Statistical analysis

All variables of interest in the study population were summarized using descriptive statistics. For continuous variable age, means and standard deviations were generated. Univariate analysis was conducted to generate frequencies and percentages for categorical variables and were used to describe the characteristics of the study population in relation to relevant variables. Proportions were compared by using Chi^2^ tests, or Fisher’s exact tests, as appropriate. McNemar χ2 test to determine the change between pre and post-intervention knowledge levels were statistically significant. The impact of socio-demographic characteristics on knowledge levels of cervical cancer was investigated using bivariate method. Binary logistic regression used to find out the statistical association between the outcome variable and the explanatory variables. Finally, explanatory variables with p-value less than 0.2 in the bivariate analysis were included and multivariate and multinomial linear regression analyses were conducted to investigate factors predict cervical cancer and Pap smear test awareness and/or utilization of Pap smear test and to examine the correlation of baseline cervical cancer knowledge scores as well as changes in scores after the educational intervention. Odds ratio and 95% confidence interval were also used to identify the presence and strength of association wherever appropriate. All tests of significance were two-tailed at 5% level. For regression analysis, the reference category was the most common category of an independent variable (IV).

## Results

A total of 283 study participants, both from biological and non-biological sciences attended the educational training on cervical cancer general awareness and responded to both pre-intervention and post-intervention questionnaires (Table 1). The dependent variables (DVs) were compared descriptively with respect to socio-demographic characteristics. The categorical variables were expressed as percentages. Pre-post education intervention differences for knowledge scores and the proportion of correct responses for each question summarized (Table 2). Baseline knowledge was low among all groups, with scores better among older participants. The baseline knowledge about awareness, symptoms, risk factors, HPV, screening and vaccination were low among non-biological science students (Table 3). A brief, structured presentation increased cervical cancer awareness knowledge among all groups. On average, knowledge scores significantly improved from 8 to 26 after the presentation (maximum possible score 42; P < .001), irrespective of region, year of study, branch of study, and age. The baseline average score of 9 for students age 20 and above and 7 in students below 20 years, and after education intervention score increased to 24 and 28 in age 20 years below and above groups respectively. Fourth-year and above students showed a baseline score of 11 and first-year students had the lowest baseline score 6 irrespective of the branch. After education intervention, the average score of students increased in the order of third year 31, fourth year 29, first year 27 and 22 second year.

**Table 1:**
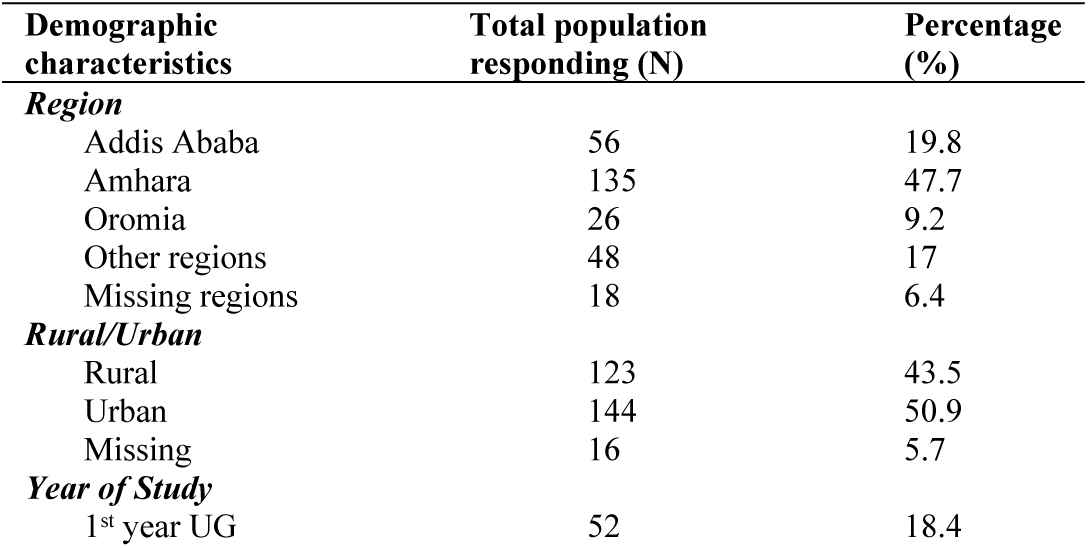
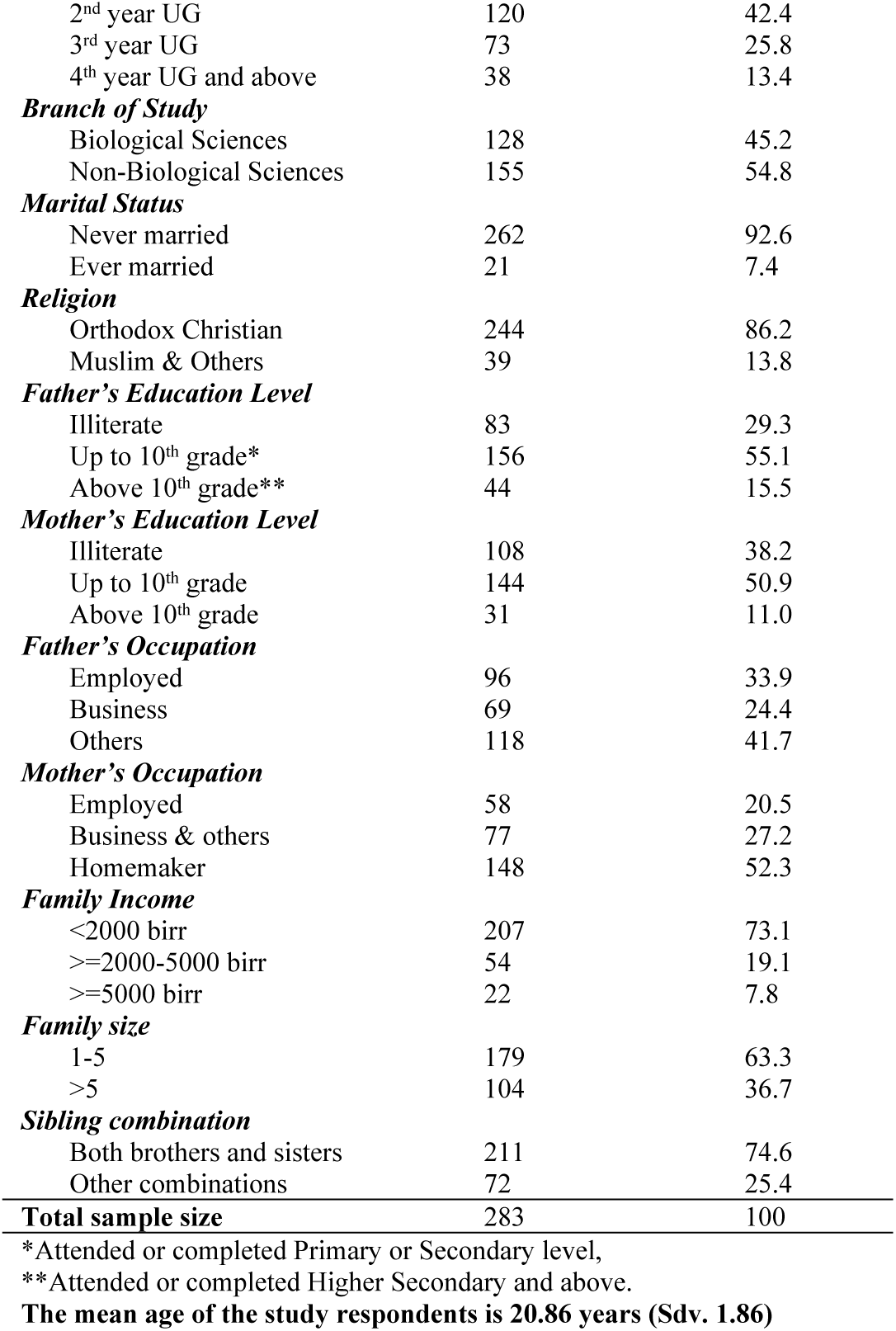
Socio-economic characteristics of study respondents.

**Table 2:** Awareness and sources of information about cervical cancer.

| Domains inquired | Pre-intervention |  | Post-intervention |  | The difference in awareness (%) |
| --- | --- | --- | --- | --- | --- |
|  | Number correctly responded (N) | Percent correctly responded (%) | Number correctly responded (N) | Percent correctly responded (%) |  |
| **General awareness** |  |  |  |  |  |
| Ever heard of cervical cancer. | 116 | 41.0 | 253 | 89.4 | 48.4 |
| Cervical cancer is a curable disease? | 38 | 13.4 | 142 | 50.2 | 36.8 |
| **Knowledge about symptoms** |  |  |  |  |  |
| Do you know the causes of cervical cancer are? | 23 | 8.1 | 217 | 76.7 | 68.6 |
| Persistent vaginal discharge that smells unpleasant. | 71 | 25.1 | 210 | 74.2 | 49.1 |
| Whether vaginal bleeding between periods could be a sign? | 43 | 15.2 | 178 | 62.9 | 47.7 |
| Do you think menorrhagia is a symptom for cervical cancer? | 52 | 18.4 | 203 | 71.7 | 53.3 |
| Vaginal bleeding after the menopause could be a sign. | 52 | 18.4 | 183 | 64.9 | 46.5 |
| Persistent pelvic pain could be a sign. | 39 | 13.8 | 182 | 64.3 | 50.5 |
| Discomfort or pain during sex could be a sign. | 64 | 22.6 | 174 | 61.5 | 38.9 |
| Vaginal bleeding during or after sex could be a sign. | 49 | 17.3 | 196 | 69.3 | 52.0 |
| **Knowledge about risk factors** |  |  |  |  |  |
| Whether poor hygiene is a risk factor? | 95 | 33.6 | 195 | 68.9 | 35.3 |
| Whether coitus at an early age is a risk factor? | 81 | 28.6 | 196 | 69.3 | 40.7 |
| Whether multiple sex partners is a risk factor? | 101 | 35.7 | 226 | 79.9 | 44.2 |
| Do you think unprotected intercourse could be a risk factor for cc? | 97 | 34.3 | 222 | 78.4 | 44.1 |
| Do you think consuming contraceptive pills could be a risk factor? | 49 | 17.3 | 179 | 63.3 | 46.0 |
| No knowledge of cervical cancer is a risk factor? | 103 | 36.4 | 194 | 68.6 | 32.2 |
| Swelling of the cervix is a risk factor? | 67 | 23.7 | 166 | 58.7 | 35.0 |
| Whether high parity is a risk factor? | 43 | 15.2 | 137 | 48.4 | 33.2 |
| Do you think smoking could be a risk factor for cervical cancer? | 55 | 19.4 | 193 | 68.2 | 48.8 |
| **Knowledge about Human Papilloma Virus (HPV)** |  |  |  |  |  |
| Causative organism of cervical cancer. | 47 | 16.6 | 222 | 78.4 | 61.8 |
| Are you aware what is HPV and its relation with cervical cancer? | 29 | 10.2 | 204 | 72.1 | 61.9 |
| HPV can infect women and can cause cervical cancer. | 59 | 20.8 | 224 | 79.1 | 58.3 |
| HPV is a sexually transmitted infection. | 43 | 15.2 | 194 | 68.6 | 53.4 |
| HPV infections are usually obvious and most infections resolve by themselves. | 36 | 12.7 | 134 | 47.3 | 34.6 |
| HPV cannot infect men. | 36 | 12.7 | 103 | 36.4 | 23.7 |
| HPV infections can cause genital warts. | 39 | 13.8 | 153 | 54.1 | 40.3 |
| HPV infections can cause oral/pharyngeal cancer. | 33 | 11.7 | 138 | 48.8 | 37.1 |
| HPV infections can cause anal cancer. | 27 | 9.5 | 59 | 20.8 | 11.3 |
| **Knowledge about screening** |  |  |  |  |  |
| Ever heard of screening. | 64 | 22.6 | 215 | 76.0 | 53.4 |
| Are you aware of any screening method? | 57 | 20.1 | 258 | 91.2 | 71.1 |
| Have you ever heard of the Pap smear test? | 84 | 29.7 | 233 | 82.3 | 52.6 |
| Pap smear test is used for. | 38 | 13.4 | 156 | 55.1 | 41.7 |
| At what age women should start screening. | 50 | 17.7 | 161 | 56.9 | 39.2 |
| How often a woman should undergo screening. | 78 | 27.6 | 175 | 61.8 | 27.6 |
| Pap smear test can pick up cell changes that may go to become cc. | 16 | 5.7 | 176 | 62.2 | 56.5 |
| **Knowledge about vaccination** |  |  |  |  |  |
| HPV vaccine exists that protects against cervical cancer. | 40 | 14.0 | 216 | 76.3 | 62.3 |
| A vaccine for HPV is not available to men. | 23 | 8.1 | 84 | 29.7 | 21.6 |
| Can HPV vaccines be given to boys? | 25 | 8.8 | 143 | 50.5 | 41.7 |
| To which age group HPV vaccine should be given. | 14 | 4.9 | 51 | 18.0 | 13.1 |
| HPV vaccine exists that can protect against genital warts. | 45 | 15.9 | 160 | 56.5 | 40.6 |
| HPV vaccines available to protect against non-cervical cancers? | 13 | 4.6 | 89 | 31.4 | 26.8 |
| Most appropriate stage for HPV vaccination. | 39 | 13.8 | 119 | 42.0 | 28.2 |
| **Total sample size** | **283** | **100** | **283** | **100** |  |

**Table 3:** Impact of education intervention on cervical cancer awareness on biological (**BS**) and non-biological science (**NBS**). **Pre IV**: Pre-intervention; **Post IV**: Post intervention. **KL**: Knowledge level; **Zero**=No knowledge; **Poor**=1-3 correct responses; **Fair**: 4-6 correct responses; **Good:** 6 and above correct responses. Values are in percentage **(%)** at P=0.05 significance level; NS=Not significant.

| Group<br>KL | Symptoms |  |  |  | Risk Factors |  |  |  | Human Papilloma Virus |  |  |  | Screening |  |  |  | Vaccination |  |  |  |
| --- | --- | --- | --- | --- | --- | --- | --- | --- | --- | --- | --- | --- | --- | --- | --- | --- | --- | --- | --- | --- |
|  | Pre IV |  | Post IV<sup>NS</sup> |  | Pre IV |  | Post IV |  | Pre IV |  | Post IV |  | Pre IV |  | Post IV<sup>NS</sup> |  | Pre IV<sup>NS</sup> |  | Post IV<sup>NS</sup> |  |
|  | BS | NBS | BS | NBS | BS | NBS | BS | NBS | BS | NBS | BS | NBS | BS | NBS | BS | NBS | BS | NBS | BS | NBS |
| **Zero** | 29.7 | 47.1 | 3.1 | 7.1 | 27.3 | 42.6 | 3.9 | 10.3 | 49.2 | 65.8 | 3.9 | 10.3 | 28.9 | 38.1 | 4.7 | 6.5 | 60.9 | 69.7 | 8.6 | 10.3 |
| **Poor** | 39.8 | 35.5 | 7.8 | 11.0 | 32.8 | 27.7 | 10.9 | 3.9 | 32.8 | 21.9 | 14.8 | 14.8 | 42.2 | 47.1 | 9.4 | 7.1 | 29.7 | 20.0 | 33.6 | 23.2 |
| **Fair** | 19.5 | 16.1 | 21.9 | 13.5 | 28.1 | 26.5 | 34.4 | 36.8 | 14.1 | 11.6 | 43.8 | 48.4 | 22.7 | 14.2 | 28.9 | 31.6 | 8.6 | 8.4 | 37.5 | 45.8 |
| **Good** | 10.9 | 1.3 | 67.2 | 68.4 | 11.7 | 3.2 | 50.8 | 49.0 | 3.9 | 0.6 | 37.5 | 26.5 | 6.3 | 0.6 | 57.0 | 54.8 | 0.8 | 1.9 | 20.3 | 20.6 |

### Socio-demographic characteristics of the study population

Demographic characteristics of the 283 female students are summarized in Table 1. Students belong to first year (18.4%), second-year (42.4%), third-year (25.8%) and fourth year & above (13.4%). The students belonged to biological sciences (45.2%) and non-biological sciences (54.8%). The mean age was 20.86 years (Sdv. 1.86) (17– 30 years) with 45.9% in 17– 20-year-old range and 54.1% in 21 and above years range. Students belong to Addis Ababa (19.8%), Amhara (47.7%), Oromia (9.2%), other regions (17%) and missing regions (6.4%) and were belong to either rural (43.5%), or urban (50.9%) and 5.7% of students’ information was missing, not included in the analysis. Majority of the participants 244 (86.2%) were Orthodox Christians, while 39 (13.8%) belonged to other religions (Muslims and other Christians). Most of the respondents 262 (92.6%) were never married and 21 (7.4%) students were married. Study participants father’s educational levels were, illiterates 83 (29.3%), up to 10^th^ grade 156 (55.1%) and above 10^th^ grade 44 (15.5%) and mother’s educational levels were, illiterate 108 (38.2%), up to 10^th^ grade 144 (50.9%) and above 10^th^ grade 31(11.0%). Respondents father’s occupation was either employed 96 (33.9%), business, 69 (24.4%) or other occupation 118 (41.7%). Only 58(20.5%) of the participant’s mothers were employed, 77 (27.2%) were either business or related occupation and most were 148 (52.3%) homemakers. 179 (63.3) had 5 or less numb of siblings and 104 (36.7%) had >5 siblings. 211 (74.6%) had both brothers and sisters and 72 (25.4%) belonged to other combinations (only brothers/sisters/no sibling). Most of the participants 207 (73.1%) family income were <2000 birr, 54 (19.1%) families had monthly income >=2000-5000 birr and only 22 (7.8%) had >=5000 birr as monthly income. Responses to questions on selected domains were presented in table 2.

### Awareness of women about cervical cancer and its preventable nature

The women were asked if they have ever heard of cervical cancer. Before education intervention, one hundred sixteen (41.0%) women reported that they had heard about cervical cancer and after education intervention two hundred and fifty-three students (89.4%) aware about cervical cancer (Table 2). 38 (13.4%) participants were well aware of the preventable nature of cervical cancer before education intervention (Table 2) of this 25 (66%) participants belonged to biological sciences and 13 (34%) were belong to non-biological sciences (Table 3). After educational intervention 142 (50%) could learn the preventable nature of cervical cancer and of this 67 (47%) participants belonged to biological sciences and 75 (53%) were belong to non-biological sciences.

### Knowledge about the symptoms of cervical cancer

Eight questions were asked about the symptoms, and at baseline knowledge was the least about the causes of cervical cancer, only 23 (8.1%) students correctly answered and but after education intervention 217 (76.7%) students reported that they know about the causes of cervical cancer (Table 2). Similarly, before the intervention, persistent vaginal discharge could be a symptom was most correctly answered by 71 (25.1%) students. However, one hundred and fifty-five (54.77%) of the respondents did not know any symptom and symptoms associated with cervical cancer before educational intervention and this includes 98 (63.22%) respondents from non-biological sciences and 57 (36.77%) biological sciences. After educational intervention, 90.46% of study respondents could respond to any of the cervical cancer symptoms correctly. Non-biological science students showed a higher increase in awareness about symptoms compared to students belong to biological sciences (Table 3). Overall mean level knowledge about the symptoms of cervical cancer before the intervention was 1.74 after education intervention was 6.81 with a mean increase of 5.07 (Table 5).

**Table 4a & 4b:**
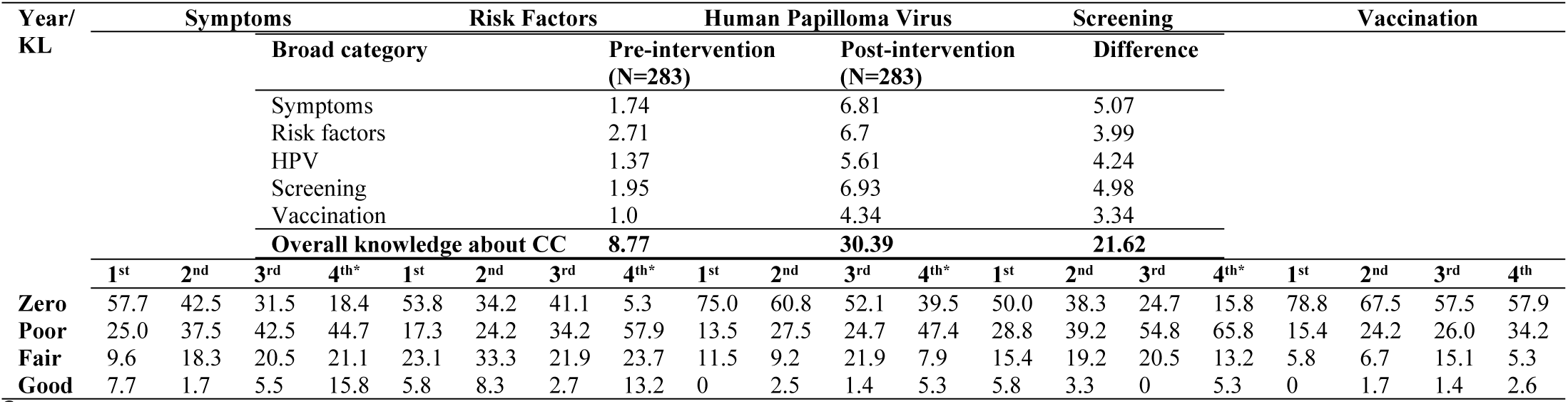
Impact of education intervention on cervical cancer awareness on the year of study. **Pre IV**: Pre-intervention; **Post IV**: Post intervention. **KL**: Knowledge level; **Zero**=No knowledge; **Poor**=1-3 correct responses; **Fair**: 4-6 correct responses; **Good:** 6 and above correct responses. Values are in percentage **(%)** at P=0.002 significance level.

**Table 5:**
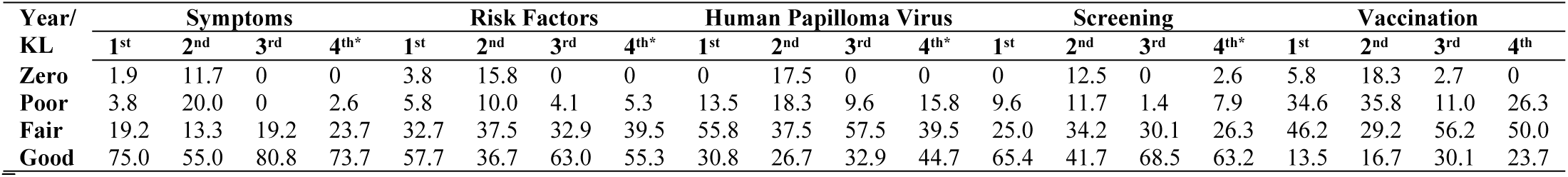
Mean level of awareness on various broad issues (categories) of Cervical Cancer (CC)

### Knowledge about the risk factors of cervical cancer

To assess knowledge about the cervical cancer risk factors, nine questions including multiple sexual partners, poor hygiene, no knowledge on cervical cancer and cigarette smoking could promote cervical cancer were asked to study participants (Table 2). About 35.6 % (n=101) study respondents had no idea about risk factors associated with the disease before educational intervention and only 7.4% (n=21) students could not identify any of the risk factors even after educational intervention. Before the intervention, 43 (15.2%) students felt high parity could be a risk factor and after the educational intervention, 137 (48.4%) could felt high parity could be a risk factor and it was the least correctly responded question among the nine risk factors were asked. One hundred and eighty-two (64.4%) study participants were able to identify minimum one risk factor before intervention and 101 (35.6%) includes 66 (22.96%) respondents from non-biological sciences and 35 (12.36%) respondents of biological sciences could not identify a single risk factor correctly. After educational intervention, biological science students showed the highest increase in awareness about risk factors compared to students from non-biological sciences (Table 3). More than 95 (33%) students identified, multiple sex partners, poor hygiene, no awareness of cervical cancer, unprotected intercourse could be risk factors (Table 2). Mean baseline awareness about the risk factors was 2.71, which was highest compared to other categories of the questionnaire and after intervention an overall increase of 6.7.

### Knowledge about the HPV and its relationship with cervical cancer

Nine different questions like the causative organism, mode of transmission of HPV and different diseases in male and females were asked about HPV and its relationship with cervical cancer to understand the knowledge levels before and after the educational intervention. Before the educational intervention, 43 (15.2%) of study respondents were not aware of STD nature of HPV infections and different diseases caused by it and 194 (68.6%) women responded correctly after post-intervention (Table 2). Before the educational intervention, 16.6% and after intervention 78.4% female students aware HPV as cervical cancer causative organism. HPV can cause anal cancers was the least correctly answered before (9.5%) and even after (20.8%) education intervention. Twenty-nine (10.2%) respondents before intervention were aware of HPV and its relationship with cervical cancer and 204 (72.1%) students identified correctly the HPV relationship with mean 61.9% increase after post-intervention (Table 2). 102 (36.04%) respondents from non-biological sciences and 62 (21.9%) biological sciences had no baseline awareness about HPV and its relationship with different diseases. After educational intervention, non-biological science students showed the highest increase in awareness about HPV compared to students belong to biological sciences (Table 3). Overall mean knowledge level before the intervention was 1.37, and after the intervention was 5.61 with an increase of 4.24 (Table 5).

### Knowledge about the screening of cervical cancer

There were seven different questions like heard of cervical cancer screening, any screening method, Pap smear test, and its importance, when should women start screening and how often should be screened. Before the educational intervention, only 19.7% of total respondents were aware of screening and 69.32% women could respond correctly after intervention (Table 2). How often women should undergo screening was correctly responded by 27.6% before intervention and 61.8% of respondents answered correctly after the intervention. Only 5.7% of respondents’ identified Pap smear test can pick cell changes before intervention and it increased to 62.2% after educational intervention. 95 (33.56%) study respondents had no baseline knowledge about screening and its importance with 37 (13.07%) respondents belong to biological and 58 (20.49%) non-biological sciences. After intervention, 43.1% (n=122) biological and 51.23% (n=145) non-biological sciences showed awareness about screening (Table 3). However, before the intervention, 11.3% each from biological and non-biological sciences were heard of cervical cancer screening and after the intervention, it was increased to 32.1% and 43.8% in biological and non-biological sciences. 8% and 5.3% before and after 39.2% and 43.1% of biological and non-biological sciences from total respondents reported that they were heard of Pap smear test. After educational intervention, increase in awareness about cervical cancer screening was good in respondents from non-biological sciences over biological sciences (Table 3). Overall mean level of knowledge before the intervention was 1.95 after the intervention was 6.93 with a mean increase of 4.98 (Table 4).

### Awareness regarding the target population for HPV vaccination

There were eight different questions like availability of HPV vaccine, the age of vaccination, availability of HPV vaccine both for girls and boys, a vaccine for non-cervical cancers were asked before and after the educational intervention. 48.5% of total respondents before and 91.5% after education intervention were aware of HPV vaccination (Table 2). HPV vaccination category was least understood even after education intervention. 65% of study participants showed no baseline knowledge about vaccine category. Baseline knowledge about two important knowledge indicators, availability of a vaccine to protect non-cervical cancer, was 4.6% and age group for vaccination was 4.8%. After the intervention, only 18% of study participants correctly understand the right age for vaccination in girls. HPV vaccines could be given to boys, 8.8% before and after intervention 50.5% (P=0.05) could respond correctly. 18% of study respondents before and 42.4% after intervention responded correctly to the best time for HPV vaccination would be before becoming sexually active. 107 (37.8%) respondents from non-biological sciences and 78 (27.56%) respondents of biological sciences showed no baseline knowledge about the vaccine and its importance. After educational intervention, non-biological science students showed the highest increase of awareness that students belong to biological sciences (Table 3). Overall mean knowledge level before the intervention was 1.0, and after the intervention was 4.34 with a mean increase of 3.34 (Table 5).

### Source of information about cervical cancer and Pap smear test

41% respondents said they heard about cervical cancer through some source before intervention and after educational intervention increased to 90%. Most common source of information at baseline was other sources (22.6%) including media. After educational intervention, health educator score was increased from 11.3 to 47.4% (Table 6). 29.7% of respondents who had heard about Pap smear test got their information from the medical staff, followed by other sources (mass media), relatives and friends (Table 6). After educational intervention, 82.6% of respondents were reported awareness of Pap smear and sources of Pap smear test. Health educator as the source of information before and after the educational intervention increased from 13% to 61.4%. Utilization of the Pap smear test only once among this population and only 3.5% of participants ‘family members’, being screened (Table 7).

**Table 6:** Source of information about cervical cancer and Pap smear test knowledge before and after education intervention and influence of the branch of study during pre-intervention (* Chi2= 9.54 & Cramer’s V=.184) (**Chi2=9.61 & Cramer’s V= .184) at the P=0.05 significance level.

| Sources of information | The number responded (N) | Percent responded (%) | The number responded (N) | Percent responded (%) | Change in response after intervention (%) |
| --- | --- | --- | --- | --- | --- |
| **Respondents source of information about cervical cancer knowledge\*** | N=116 | %=41.0 | N=253 | %=90 |  |
| Parents/ Family Members/Relatives | 13 | 4.6 | 21 | 7.4 | 2.8 |
| Medical or nursing staff/Health Educator/Teacher | 32 | 11.3 | 134 | 47.4 | 36.1 |
| Friends/Classmates | 7 | 2.5 | 17 | 6.0 | 3.5 |
| Other sources | 64 | 22.6 | 82 | 29.0 | 6.4 |
| **Respondents source of information about Pap smear test\*\*** | N=84 | %=29.7 | N=235 | %=82.6 |  |
| Parents/ Family Members/Relatives | 15 | 5.3 | 15 | 5.3 | 0.0 |
| Medical or nursing staff/Health Educator/Teacher | 37 | 13.0 | 175 | 61.4 | 48.4 |
| Friends/Classmates | 8 | 2.8 | 21 | 7.4 | 4.6 |
| Other sources | 24 | 8.5 | 24 | 8.5 | 0.0 |

**Table 7:**
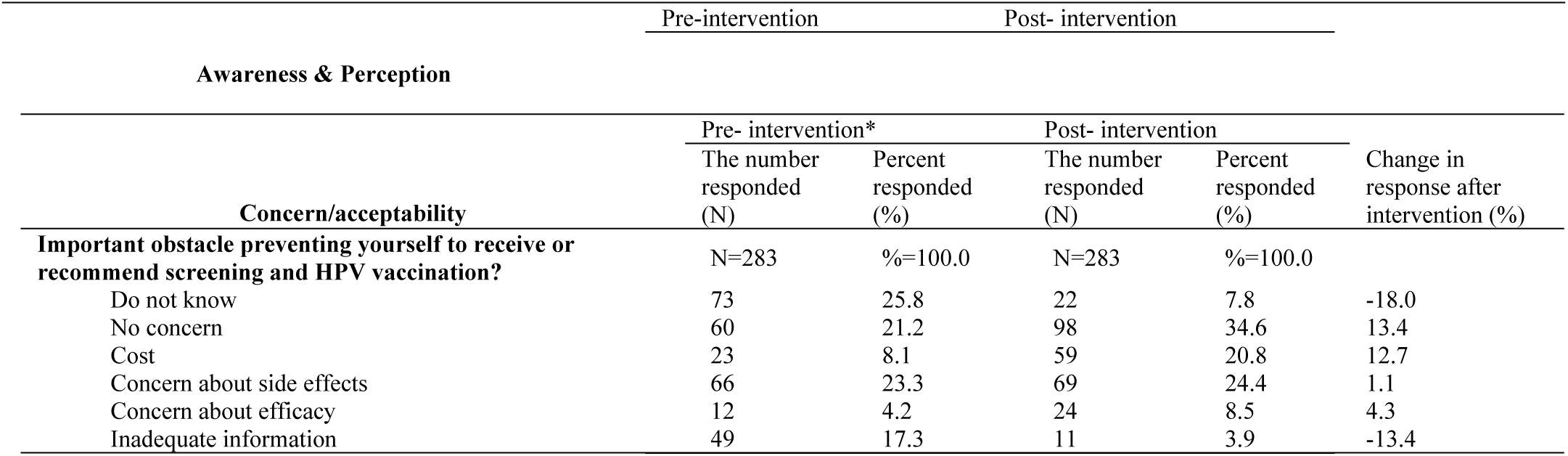
Impact of education intervention on awareness about Pap smear test was done and perceptions to receive cervical cancer screening and vaccination.

### Perceptions of cervical cancer screening and HPV vaccination

Perceptions of cervical cancer screening and HPV vaccination of the respondents are presented in Table 7. Before the educational intervention, 25.8% and after 46.3% of the respondents would like to receive or recommend cervical cancer screening. Similarly, 15.9% of respondents before the educational intervention, 47% of the respondents after educational intervention would like to receive or recommended for HPV vaccination. Before the educational intervention, 9.5% biological and 6.3% non-biological sciences expressed acceptance for HPV vaccination and after the intervention, acceptance was increased to 18.3% and 28.2% respectively. From total respondents, 1% of first year, 9.8% of second year, 1.4% of third year and 3.5% of fourth-year students expressed likeliness to receive HPV vaccination before intervention and after educational intervention, 10.6% (first year), 16.6% (second year), 14.8% (third year) and 4.9% (fourth year & above) students agreed (Table 4).

### Concerns of receiving or recommending HPV vaccination

Overall acceptance of HPV vaccine among the study population before 21.2% and 34.6% after educational intervention. Before and after educational intervention concern about side effects (23.3%, 24.4%), efficacy (4.2%, 8.5%), inadequate information (17.3%, 3.9%), and cost (8%, 20.8%) respectively. Interesting inadequate information as a complaint reduced from 17.3% to 3.9% (Table 8).

**Table 8:**
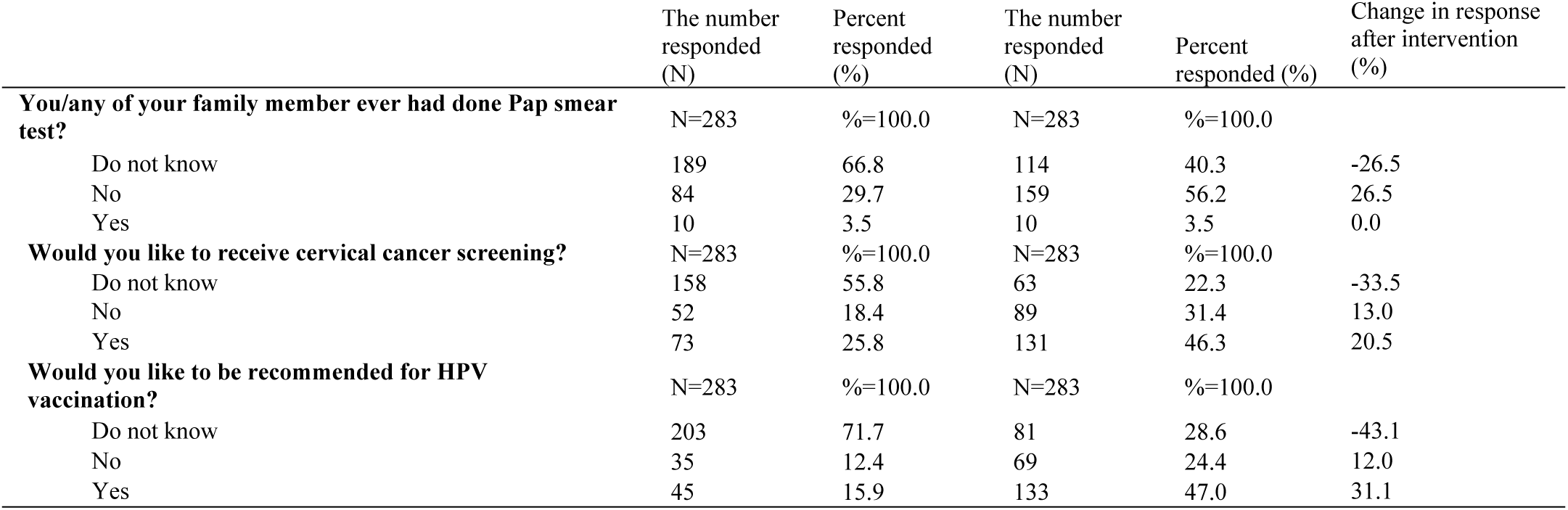
Concerns/acceptability to take up cervical cancer screening before and after education intervention and influence of *Age category at pre-intervention with Chi2= 15.90 at the P=0.05 significance level.

### Health seeking behaviour of respondents before and after education intervention

To understand the health-seeking behaviour, respondents were been asked if they have a symptom of cervical cancer, how soon they visit a doctor and in response to this, 58% of respondents could not decide before intervention and 25.8% could not understand the importance of health check even after education intervention. 1.4% of respondents before and 9.9% after intervention said, they never visit any medical help. Before intervention, 23.7% and after intervention, 39.2% respondents reported, they will visit medical hospital within a few days. 18.3% before and 34.9% after education intervention felt they will visit hospital from a couple of weeks to a couple of months (Table 9).

**Table 9:**
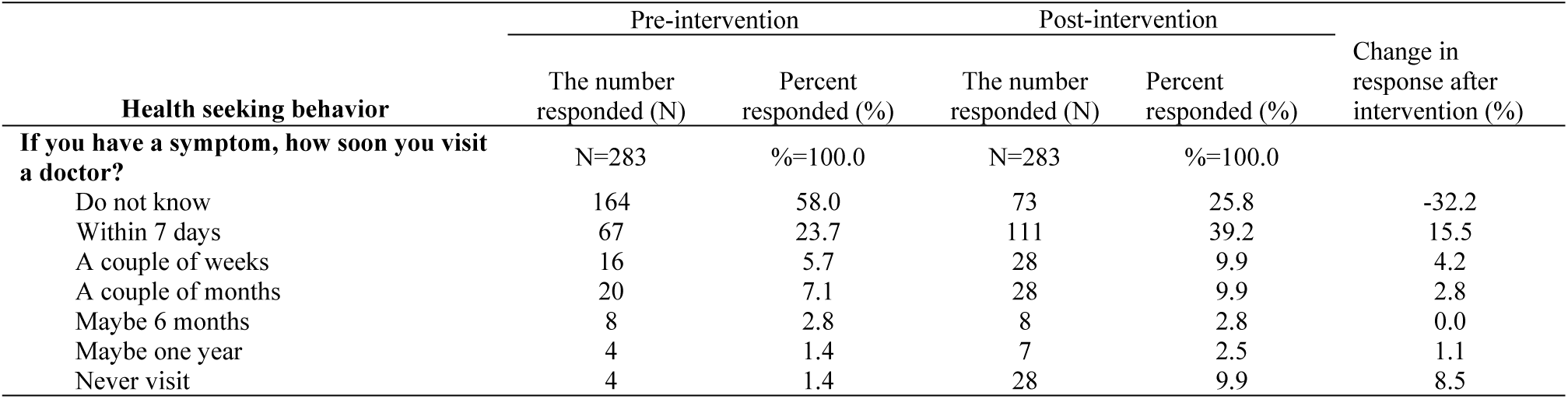
Health seeking behavior of respondents before and after education intervention and influence of the branch of study at post-intervention with Chi2= 31.81 and Cramer’s V= 0.335 at the P=0.05 significance level.

### Preference of venue for cervical cancer screening and vaccination before and after education intervention

Before the educational intervention, 53% and after the intervention, 30% of the respondents could not decide the preference of venue for the screening and vaccination. Before the intervention, women and children’s hospital was the most preferred venue (14.5%) and after the educational intervention, local community health centre/local clinic was the preferred venue (28.3%). General hospital as venue preferred by 9.2% respondents before and after intervention (Table 10).

**Table 10:** Preference of venue for cervical cancer screening and vaccination before and after education intervention and influence of age category at post-intervention with Chi2= 10.25 and Cramer’s V= 0.190 at the P=0.05 significance level.

| Venue | Pre- intervention |  | Post- intervention |  | Change in response after intervention (%) |
| --- | --- | --- | --- | --- | --- |
|  | The number responded (N) | Percent responded (%) | The number responded (N) | Percent responded (%) |  |
| **The appropriate venue for screening and vaccination** | N=283 | %=100.0 | N=283 | %=100.0 |  |
| Do not know | 150 | 53.0 | 85 | 30.0 | -23.0 |
| Local Community health center/clinic | 39 | 13.8 | 80 | 28.3 | 14.5 |
| Women and children's hospital | 41 | 14.5 | 56 | 19.8 | 5.3 |
| General hospital | 26 | 9.2 | 26 | 9.2 | 0.0 |

| Factors | Cervical cancer screening |  |  |  | Cervical cancer vaccination |  |  |  |
| --- | --- | --- | --- | --- | --- | --- | --- | --- |
|  | Pre-intervention |  | Post-intervention |  | Pre-intervention |  | Post-intervention |  |
| | $\chi^2$ | Cramer's V | $\chi^2$ | Cramer's V | $\chi^2$ | Cramer's V | $\chi^2$ | Cramer's V |
| Age category | 5.984 | .145 | 9.456 | .183 | 10.19 | .190 | 6.66 | .154 |
| Year of study | 16.45 | .171 | 36.94 | .255 |  |  | 32.21 | .239 |
| Branch of study |  |  | 11.05 | .198 | 10.02 | .188 |  |  |
| Religion |  |  | 7.64 | .164 |  |  | 7.02 | .158 |
| Father's education level |  |  | 14.88 | .162 |  |  | 11.01 | .140 |
| Mother's education level | 7.29 | .113 | 12.92 | .151 |  |  | 19.24 | .184 |
| Father's occupation | 19.18 | .184 | 13.86 | .157 | 11.93 | .145 |  |  |
| Family Income |  |  |  |  |  |  | 16.78 | .172 |
| SIB combination |  |  |  |  | 8.80 | .176 |  |  |
| School |  |  | 27 | 9.5 | 36 | 12.7 |  | 3.2 |

### Overall knowledge about cervical cancer and associated factors

#### Chi^2^ test of independence and McNemar’s test

To understand the influence of various socio-demographic factors on perception of cervical cancer screening and HPV vaccination, a Chi^2^ test of independence was carried out. Age and father’s occupation had a significant impact on both screening and vaccination before educational intervention. The post-intervention perception was under the influence of age, year of study, religion, parents’ education level, and family income at P=0.05 (Table 11). Bivariate analysis showed six socio-demographic characteristics were found to be significantly associated with knowledge levels about cervical cancer: age, educational level, branch of study, fathers and mother’s education levels, and family size (Table 12).

**Table 11:** Factors influencing perceptions to receive cervical cancer screening and vaccination before and after educational intervention at P=0.05.

Chi<sup>2</sup> test of independence and McNemar's test
| Variables checked for association | Age |  | BOS |  | YOS | F. Ed. | M. Ed. | FS |
| --- | --- | --- | --- | --- | --- | --- | --- | --- |
|  | Pre | Post | Pre | Post | Post | Pre | Pre | Post |
| **General awareness and symptoms** |  |  |  |  |  |  |  |  |
| Ever heard of cervical cancer |  | 7.9 | 7.845 | 9.3 | 14.3 |  |  | 7.48 |
| Cervical cancer is a curable disease? | 8.75 |  | 7.49 |  | 10.3 |  |  |  |
| Do you know the causes of cervical cancer are? |  |  | 11.02 |  | 34.9 |  |  | 6.34 |
| Persistent vaginal discharge that smells unpleasant. | 5.61 |  | 5.99 |  | 29.12 |  |  | 6.54 |
| Whether vaginal bleeding between periods could be a sign? |  |  | 4.75 |  | 16.7 |  |  |  |
| Do you think menorrhagia is a symptom for cervical cancer? |  |  | 3.99 |  | 17.0 |  | 13.55 |  |
| Vaginal bleeding after the menopause could be a sign. | 4.49 | 4.0 |  | 5.37 |  |  | 15.02 | 17.37 |
| Persistent pelvic pain could be a sign. |  |  | 4.85 |  | 14.74 |  | 6.85 | 10.87 |
| Discomfort or pain during sex could be a sign. |  |  | 4.05 |  |  | 7.18 | 7.41 | 9.0 |
| Vaginal bleeding during or after sex could be a sign. | 5.6 |  | 4.65 |  | 10.8 |  |  | 7.0 |
| **Knowledge about risk factors** |  |  |  |  |  |  |  |  |
| Whether poor hygiene is a risk factor? | 5.92 | 4.87 |  |  |  | 8.97 |  |  |
| Whether coitus at an early age is a risk factor? |  |  |  | 5.93 |  |  |  | 5.73 |
| Whether multiple sex partners is a risk factor? |  |  | 12.74 | 4.16 | 12.0 |  | 7.87 | 4.58 |
| Do you think unprotected intercourse could be a risk factor? |  |  |  |  | 19.5 | 8.11 | 8.11 | 3.8 |
| Do you think consuming contraceptive pills could be a risk factor? |  |  |  |  | 9.48 |  | 6.64 | 3.9 |
| No knowledge of cervical cancer is a risk factor? |  | 4.34 | 6.68 |  | 15.28 | 9.69 | 16.74 | 7.35 |
| Swelling of the cervix is a risk factor? |  | 3.99 | 5.97 |  |  | 9.84 |  | 7.55 |
| Whether high parity is a risk factor? |  | 9.59 |  |  | 28.2 |  |  | 7.9 |
| Do you think smoking could be a risk factor for c. cancer? |  |  |  |  | 30.5 |  |  | 4.34 |
| **Knowledge about Human Papilloma Virus** |  |  |  |  |  |  |  |  |
| Causative organism of cervical cancer. |  |  | 16.7 | 11.8 | 34.45 |  |  | 6.45 |
| Human papillomavirus (HPV) and its relation to cc? | 4.38 |  |  |  | 28.62 | 6.9 | 6.1 |  |
| HPV can infect women and can cause cervical cancer. |  | 6.0 | 7.4 |  | 31.5 |  |  |  |
| HPV is a sexually transmitted infection. |  |  | 4.7 | 5.74 |  | 6.0 |  |  |
| HPV infections are usually obvious and most infections resolve by themselves. | 9.34 | 4.1 | 5.0 |  |  |  |  |  |
| HPV cannot infect men. |  |  |  | 9.5 | 13.4 |  |  |  |
| HPV infections can cause genital warts. | 5.94 | 8.67 | 6.5 |  | 20.3 | 9.43 |  |  |
| HPV infections can cause oral/pharyngeal cancer. | 5.46 |  |  |  | 15.26 |  |  |  |
| HPV infections can cause anal cancer. |  | 4.42 |  | 13.1 | 16.16 |  |  | 5.7 |
| **Knowledge about screening** |  |  |  |  |  |  |  |  |
| Ever heard of screening. |  | 4.69 |  |  | 22.8 |  | 11.9 |  |

**Table 12:**
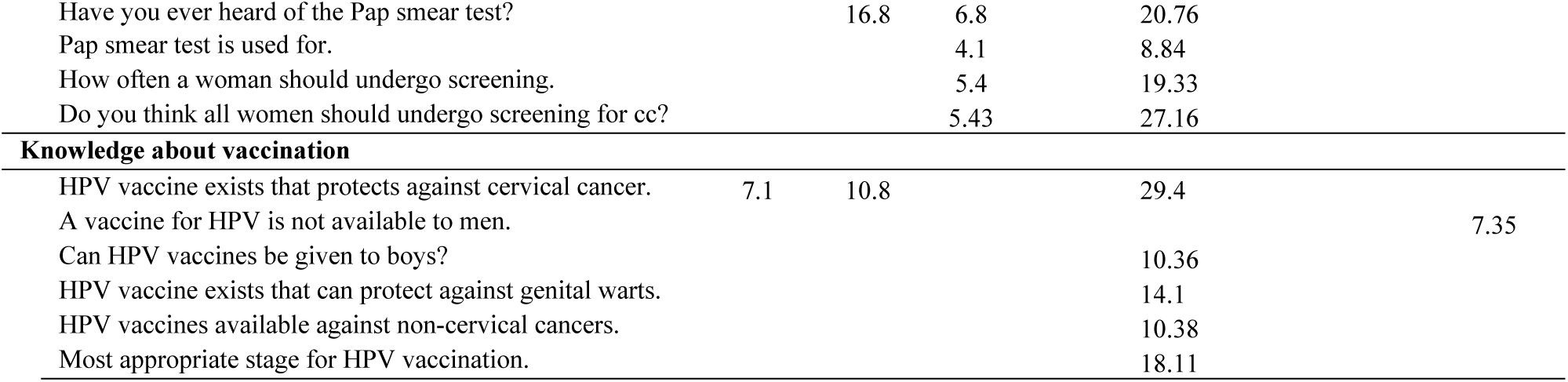
Chi-square analysis of independence of various socio-demographic factors and dependable variables about cervical cancer symptoms, Risk factors, HPV, screening and vaccination of respondents during pre and post educational intervention at P=0.05 significance. BOS= Branch of study; YOS=Year of study; FE=Father’s education level; ME= Mother’s education level; FS=Family size.

**Table 13:** McNemar test of cervical cancer awareness about symptoms, risk factors, HPV, screening and vaccination of respondents at P=0.000 significance level.

| Domains compared | N= 283<br>Chi<sup>2</sup> value |
| --- | --- |
| **General awareness** |  |
| Ever heard of cervical cancer | 131.177 |
| Cervical cancer is a curable disease? | 84.198 |
| Do you know the causes of cervical cancer are? | 190.046 |
| Persistent vaginal discharge that smells unpleasant. | 116.834 |
| Whether vaginal bleeding between periods could be a sign? | 114.369 |
| Do you think menorrhagia is a symptom for cervical cancer? | 139.752 |
| Vaginal bleeding after the menopause could be a sign. | 119.858 |
| Persistent pelvic pain could be a sign. | 133.536 |
| Discomfort or pain during sex could be a sign. | 86.094 |
| Vaginal bleeding during or after sex could be a sign. | 126.130 |
| **Knowledge about risk factors** |  |
| Whether poor hygiene is a risk factor? | 71.022 |
| Whether coitus at an early age is a risk factor? | 88.408 |
| Whether multiple sex partners is a risk factor? | 101.828 |
| Do you think unprotected intercourse could be a risk factor for cervical cancer? | 104.599 |
| Do you think consuming contraceptive pills could be a risk factor for cervical cancer? | 115.563 |
| No knowledge of cervical cancer is a risk factor? | 63.780 |
| Swelling of the cervix is a risk factor? | 75.622 |
| Whether high parity is a risk factor? | 64.545 |
| Do you think smoking could be a risk factor for cervical cancer? | 118.791 |
| **Knowledge about Human Papilloma Virus** |  |
| Causative organism of cervical cancer | 165.443 |
| Are you aware what is Human papillomavirus (HPV) and its relation with cervical cancer? | 165.443 |
| HPV can infect women and can cause cervical cancer | 127.592 |
| HPV is a sexually transmitted infection | 131.579 |
| HPV infections are usually obvious and most infections resolve by themselves | 71.280 |
| HPV cannot infect men | 44.000 |
| HPV infections can cause genital warts | 95.291 |
| HPV infections can cause oral/pharyngeal cancer | 82.565 |
| HPV infections can cause anal cancer | 12.645 |
| **Knowledge about screening** |  |
| Ever heard of screening | 133.136 |
| When should women start screening? | 84.615 |
| Have you ever heard of Pap smear test | 141.316 |
| Pap smear test is used for. | 103.705 |
| Pap smear test can pick up cell changes that may go on to become cervical cancer. | 154.152 |
| How often should women have cervical cancer screening? | 111.455 |
| **Knowledge about vaccination** |  |
| HPV vaccine exists that protects against cervical cancer. | 154.230 |
| A vaccine for HPV is not available to men | 40.449 |
| Can HPV vaccines be given to boys? | 96.401 |
| To which age group HPV vaccine should be given | 21.966 |
| HPV vaccine exists that can protect against genital warts | 88.408 |
| HPV vaccines available that protect against non-cervical cancers | 62.500 |
| Most appropriate stage for HPV vaccination | 63.366 |

Age, branch of study, father’s and mother’s education level had strong association on awareness before intervention (Table 12) and post-intervention knowledge gain was under the strong influence of year of study and other influencing factors were age, branch of study, family size. Age, educational level and branch of the study were found to have a significant association with level of knowledge about cervical cancer before and after intervention (Table 12). McNemar test of cervical cancer awareness was carried out and change of overall score of symptoms, risk factors, HPV, screening and vaccination of respondents was at P=0.000 significance level (Table13). Using the sum of all knowledge items, we determined that a total of 33.9% (P=0.001) of the participants had sufficient (very good) knowledge about cervical cancer after the educational intervention.

### Multi-variate statistical analysis

A multivariate analysis was done using multiple logistic regression models to investigate the predictors of awareness of symptoms, risk factors, HPV, screening, and vaccination in the study population. The result of the analysis showed that before the educational intervention, the branch of study, and after educational intervention year of study significantly predict levels of awareness of cervical cancer.

## Discussion

The main objective of this study was to assess knowledge levels at baseline and after education intervention about cervical cancer symptoms, risk factors, HPV and its relation to cervical cancer, screening, and vaccination, and factors influence the knowledge levels, this is the first kind of study carried out using questionnaire validated through pilot study to understand the impact of knowledge intervention on young 17 to 30 years aged college attending women of University of Gondar, Northwest Ethiopia region. To prevent and control any disease, knowledge is prerequisite and attitude plays a crucial role and our study showed very poor knowledge levels, similar observations from various regions of Ethiopia [4, 5, 28] and different African countries [39–45]. The baseline awareness about knowledge, symptoms, risk factors, HPV, screening and vaccination was low before intervention (18.27%) and is very lower compared to different studies from Nigeria (23.4%), Addis Ababa (34.2 %), Ghana (37.0%) and, South Ethiopia (46.3%) and 51%, Dessie town [44–48]. Developing countries have poor knowledge [49–52] compared to developed countries [53, 54]. 41.0% of our study participants heard about cervical cancer before intervention was similar with 40.8% in Nigeria [55]. However lower than reports from different regions of Ethiopia, 53.11% in Mizan Tepi, 76.8% in Hawassa, 78.7% in Gondar town [2, 29, 56] and in some African countries like Republic of Congo (81.9%), in Botswana (77%) and 68.4% in Southern Ghana [57–59]. Students of fourth year and above showed baseline score of 11 compared to first-year students (6) irrespective of the branch and can be compared with earlier study on Hawassa university students [2]. Studies show that, levels of education was significantly associated with knowledge about cervical cancer [48, 57, 60].

The baseline knowledge of biological sciences was 10 and non-biological sciences participants showed 7 and background of biological sciences might influence baseline knowledge and similar observation was reported that knowledge of medical students was better over public health students [2]. 49% of our participants’ baseline level on various symptoms associated with cervical cancer such as vaginal bleeding between periods (15.2%), painful coitus (22.6%) and bleeding after intercourse (17.3%) were reported and these findings are lower than studies carried out [61–64]. Before the educational intervention, study participants showed poor knowledge about cervical cancer risk factors. About 35.6 % of student respondents had no idea about risk factors associated with the disease before educational intervention which was very lower than 67.9% reported for Hawassa University College students [2]. 30.1% study respondents identified one or more correct risk factors before education intervention which was very much matched with study carried out at Gondar, 31% [5] and was much lower than the similar study done in South Africa, 64.0% [65]. 33% of our students identified, multiple sex partners, and similar response observed in the previous reports [66–69], however, response is higher than the study conducted in South Africa (26%), however, is lower than 49.7% awareness showed by Hawassa university medical students [2] and 53% awareness by university students of Bhutan [70] and other studies [63, 71, 72]. The difference could be due to the inaccessibility of the cervical cancer screening service, as well as less attention was given to reproductive health in Ethiopia. A study from Malaysia could not identify any of the cervical cancer risk factors [69].

Baseline knowledge about prolonged use of contraceptive pills as a risk factor was low in our study participants and only 17.3% study respondents identified and a similar observation was reported [73]. Only 19.4% of our study respondents identified smoking is a risk factor which is lower (22.3%) than a study in Gabon [74] but higher than a study in Ghana, only 1% participants identified [75]. 28.6% study respondents identified early coitus could be a risk factor and is higher than 13%, reported in a study [67]. 16.6% of the participants were aware HPV as the causative organism and is better over 9% reported in Southern Ethiopia [76] and 8% in Gabon [74] and [60]. However, is much lower than similar studies carried out in Northern Ethiopia [5] and other regions of Africa [77, 78]. Only 15.2% study respondents identified, STI nature of HPV before educational intervention and was matching with a study [79], however, was low than other reports, 31.5% [80] and 41% [67].

Low levels of knowledge on HPV was reported in a study US [81] and in another report, 78.5% of the college women to have heard of HPV in the US [82], UK, 63% [83]. Several studies from different countries reported that overall, the general public has low-level of awareness about HPV infection [84]. Only 9.5% of study respondents identified, HPV can cause anal cancers before the educational intervention and is less than similar earlier reports [85–90]. 13.4% of our participants were aware of the preventable nature of cervical cancer before education intervention and is matching with similar studies from semi-urban India, 12.2% and 11% [67, 91] and is lower than similar studies reported in 17.5%, [92], 30.5% in Burkina Faso [93], Addis Ababa, 50.6% [34], 51.5% [94](Awodele et al., 2011), South Africa, 57.0% [65], Southern Ethiopia, 57.6% [76], Northern part of Ethiopia, 63.9% [5]. Base level knowledge of 19.7% of study respondents were aware of screening, higher than 11% [67] and Malaysian population [69] and lower than 33.97% in Mizan Tepi, Ethiopia [56] and 41% [95]. A study in Addis Ababa revealed, that the vast majority of nurses and midwives had poor knowledge on aetiology and risk factors and never heard of any screening methods other than the Pap smear [12]. Only 3% of utilization of Pap smear test once among our study participants relatives and is matching with similar studies, only 5% respondents underwent Pap test [61, 96]. The low levels of awareness could be due to lack of nationwide screening policy in Ethiopia. This could be due to low levels of knowledge on cervical cancer and is supported by earlier report that, cervical cancer knowledge levels determine the rate of screening uptake [41]. This highlights the need of spread about awareness and health education about cervical cancer is critical as primary care taken to scale up the screening in Ethiopia. According to FMoH, 2016 [27], cervical cancer screening and prevention strategies are initiated by the Ethiopian government.

Before intervention, 39.5% of total respondents were not aware of HPV vaccination and 65% of study participants showed no baseline knowledge about vaccine category. 4.6% of respondents know that vaccine is available to protect against non-cervical cancer and 4.9% of respondents only aware at what age vaccine should be given, 8.8% of students answered HPV vaccines could be given to boys. Similarly, 15.8% of participants reported that vaccination could be a preventive method [76]. Most countries in sub-Saharan Africa, including Ethiopia, did not include routine HPV vaccination in the national prevention strategy for cervical cancer and other HPV-related diseases [28]. Despite vaccination, not being implemented in Ethiopia, the awareness and knowledge of participants would help as an effective primary prevention strategy [97–99]. The Ethiopian government has also recently introduced HPV vaccination demonstration project and yet to available as a national program [34]. 41% respondents said they heard about cervical cancer through some source. 29.7% of respondents from the medical staff, followed by 22.6% other sources including relatives and friends and 11.3% teachers. Similar observations reported by various studies, 55.5% teachers as the source, 30.5% mass media and 22.9% health worker as their source of information for cervical cancer and its screening [2, 100]. 29.7% of respondents who had heard about Pap smear test got their information from the medical staff, followed by other sources, relatives, and friends. This is higher than a similar type of studies carried in other places like Nigeria [44] 27%, Gondar town, 13.7% [5], not aware of the Pap smear test [101–104]. But lower than South Africa where 49.0% of the respondents heard of the test [65]. The low participation of health workers indicate that health workers are not thoroughly trained and media is not able to reach both rural and urban parts of the country equally. Women from urban areas were obtained information through various sources including, internet and mass media [105]. In contrast, a report on Congo women showed that conversation with other people was the basic source of cervical cancer awareness than through media [57]. Role of audio-visual means of spreading awareness had a mixed impact in African countries [106], remains a potentially important method of health promotion in rural low-educated communities.

Before educational intervention, 25.8% of the respondents would like to receive and recommend cervical cancer screening and similarly, 15.9% of respondents would like to receive and/or recommend HPV vaccination and is little higher than Southern Ethiopia, 14.2% [76], however, is very less than Ruvuma 55.7%, [75], Mizan Tepi University students, 61.24% [56]. An important observation in our study participants consistent with studies carried out in other African countries is, willingness for the cervical cancer screening was found poor even after having knowledge of the disease and its importance [56] and similar findings in other parts of Ethiopia [29, 77, 107]. This could be due to lower attention to female health in Ethiopia. Overall acceptance of HPV vaccine among the study population was 21.2%. The main concerns were about side effects (23.3%), efficacy (4.2%), inadequate information (17.3%), and cost (8%). Similar reports, the cost was a major concern [108, 109] and inadequate information [110] was reported. There was a low acceptance to seek the medical help in our study participants and 39.2% respondents reported, they will visit the medical hospital within a few days. This was less than 55.3%, Mizan Tepi [2], Addis Ababa [100], 1.4% of respondents said they never visit any medical help. This low acceptance to seek medical help might be due to psychological and socioeconomic reasons.

Before the educational intervention, the branch of study, and after educational intervention year of study significantly predict the level of awareness of cervical cancer. Similar observations reported [4] and reports suggest the income level also effect knowledge on cervical cancer, women with high-income level have more knowledge than women with low income. However other socio-demographic factors were not found to be statistically associated with knowledge levels [76] and not consistent with a study on Gondar community [29]. After educational intervention, an increase from 20.1% to 91.2% of study participants heard about the any of the screening methods, similarly an increase from 29.7% to 82.3% participants said they know Pap test as a screening method for early detection of cervical cancer. In our study population, baseline knowledge was low among all groups and similar observations in other studies [111], and low even in healthcare workers and physicians [85] and medium to low in teachers [66]. This indicates significant knowledge gaps in different populations globally and gaps are common [112].

Baseline knowledge of HPV was high in biological sciences and it could be the positive influence of the branch of study and it can be comparable to similar observation reported in teaching population [113]. In our study, after the educational intervention, the non-biological students’ knowledge levels improved over biological sciences. A similar finding observed in a study where knowledge level improved in health workers and was similar to those of physicians [85] after intervention. Our study participants were mostly from rural areas and deprived of mass media and this could be one of the reasons for poor baseline knowledge levels. Similar observation reported [114]. A brief, structured presentation increased cervical cancer awareness knowledge among all groups and is consistent with previous studies [115–117]. On average, knowledge scores significantly improved from 8 to 26 after the presentation (maximum possible score 42; P <0.001), irrespective of region, year of study, branch of study, and age. Recent years several studies reported a significant increase in cervical cancer knowledge in women after educational intervention [118–120].

Before education intervention, 41.0% of women reported that they had heard about cervical cancer and after education intervention 89.4% aware of cervical cancer. 13.4% of our participants were well aware of the preventable nature of cervical cancer before education intervention. Similarly reported that cervical cancer can be cured if it is diagnosed at an early stage [73]. After educational intervention, it increased to 50% and similar to a study [72], however, is lower than 84.2% observed in another study [92]. Only 50% awareness after intervention highlights the need and importance of education on cervical cancer, a similar observation reported [111]. Baseline knowledge about the causes of cervical cancer was 8.1% and after education intervention, awareness increased to 76.7% and similarly, before the intervention, 25.1% study respondents identified persistent vaginal discharge could be a symptom and increased to 74.2%. Overall mean level knowledge about the symptoms of cervical cancer after education intervention was increased from 1.74 to 6.81 with a mean increase of 5.07. After educational intervention, 92.6% of our students could identify at least one risk factor and knowledge levels on risk factors improved [67, 92, 121]. Before the intervention, 15.2% students felt high parity could be a risk factor and after the educational intervention, 48.4% could agree, high parity could be a risk factor. And in a study, 44% responded high parity as a risk factor [122]. Similar reports on parity in previous studies in Africa show high parity as a risk factor was underreported in sub-Saharan Africa [73], no report on parity [123]. In our study, most of the students had experience of high parity in their families and experienced self-serving bias and similar observations in other parts of the world [124–126]. Mean baseline awareness about the risk factors was 2.71 and after intervention an overall increase of 6.7. After the educational intervention, awareness about STD nature of HPV infections increased from 15.2% to 68.6% and similarly awareness about HPV as cervical cancer causative organism increased from baseline level 16.6% to 78.4%. HPV can cause anal cancers was the least correctly answered before (9.5%) and after (20.8%) the educational intervention. Overall mean knowledge level before the intervention was 1.37, and after the intervention was 5.61 with an increase of 4.24.

After the intervention, an increase from 19.7% to 69.32% in study respondents about cervical screening. Similarly in other studies, knowledge levels about symptoms, HPV, preventive methods improved after educational intervention [92]. 13.3% respondents before and 82.3% after intervention reported that they were heard of Pap smear test and similar observations earlier reported [127–129]. Pap smear test can pick cell changes knowledge levels increased from 5.7% to 62.2% in study participants after the educational intervention and educational intervention improved knowledge about HPV and cervical cancer screening. Similar observations reported in earlier studies [130–132]. The overall mean level of knowledge before the intervention was 1.95 after the intervention was 6.93 with a mean increase of 4.98.

48.5% of total respondents before and 91.5% after the educational intervention were aware of HPV vaccination. There is a wide global variation on cervical cancer awareness and HPV vaccination acceptability is reported in several reports [133–135]. In our study participants, HPV vaccination was the least improved category even after education intervention. The global concept of HPV vaccine is relatively new, may face challenges for general public acceptance. In our study, baseline knowledge was least in HPV vaccination and 65% of study participants not aware about HPV vaccination. But it is differing to the earlier reports, study participants showed poor knowledge on risk factors [67, 136]. After educational intervention, knowledge levels improved and vaccine acceptability increased from 15.9% to 47%. However, higher HPV vaccine acceptability reported in other studies, 70% [136], 80% [133], from 80% to 89% [138], 73% to 82% [134], high levels [139] and positive impact of educational intervention on HPV vaccine acceptance. [140–143].

After the intervention, only 18% of study participants correctly understand the right age for vaccination in girls. HPV vaccines could be given to boys, was increased from baseline knowledge of 8.8% to 50.5% (P=0.05) after the educational intervention. 18% of study respondents before and 42.4% after intervention responded correctly the best time for HPV vaccination would be before becoming sexually active. Similar observations in other studies, [144], increase incorrect response to 72.5% [138]. HPV vaccination overall mean knowledge level before the intervention was 1.0, and after the intervention was 4.34 with a mean increase of 3.34. Health educator as the source of information about Pap test before and after the educational intervention increased from 13% to 61.4%. This is differing from earlier reports on Nigeria where friends were the most important source before and after intervention [145, 146]. Friends and relatives were important source of information about cervical cancer and was corroborated by another study in Lagos which had similar findings [146]. After the educational intervention, screening acceptance levels increased from 25.8% and after 46.3% and similar increase to receive or recommend HPV vaccination from 15.9% to 47% in our study participants. Overall acceptance of HPV vaccine among the study population before 21.2% and 34.6% after educational intervention. The low increase in HPV vaccination awareness supported by other studies, inadequate knowledge of vaccine reported even in physicians, medical students, and other healthcare workers [85]. In the African continent, secondary prevention (54.6%) emphasized over primary prevention and vaccination was 23.4% [147]. Studies show after the educational intervention, knowledge on vaccination was low [113] but acceptability was high [113, 148].

Before and after educational intervention concern about side effects (23.3%, 24.4%), efficacy (4.2%, 8.5%), inadequate information (17.3%, 3.9%), and cost (8%, 20.8%) respectively. Interesting inadequate information as a complaint reduced from 17.3% to 3.9%. Similar reports on concern about effectiveness and side effects of the HPV vaccine [149]. Our study respondents’ health-seeking behaviour is not fully positive and 1.4% of respondents before and 9.9% after intervention said, they never visit any medical help and 25.8% could not even understand the importance of health check-up. However, there was an increase from 23.7% and 39.2% of respondents reported, they will visit medical hospital within a few days. Other similar reports show after educational intervention, positive attitude to uptake screening and vaccination [115, 129, 150–159].

Bivariate analysis showed age, branch of study, father’s and mother’s education level had strong association on awareness before intervention and post-intervention knowledge gain was under the strong influence of year of study and other influencing factors were age, branch of study, family size. A similar studies showed a significant impact of level of education, income [160] with awareness and knowledge on risk factors and vaccination [161]. Various studies show that independent variables like age, branch of study, level of education, parents’ education and occupation are good predictors of good knowledge levels of cervical cancer. Age, educational level and branch of the study were found to have a significant association with the level of knowledge about cervical cancer before and after the intervention. The similar report showed science students had better knowledge of HPV over students, not from a science background [67]. The impact of the educational intervention and an increase in the awareness about cervical cancer highlights knowledge dissemination continues to be an important tool in public health primary prevention. There is an urgent necessity to promote knowledge on risk factors of female cancers should reach all women, as well as men, and provide health education and community-based interventions. Such efforts could promote a positive attitude towards treatment options, outcomes, and survivorship in female cancers and improve practices could help overcome poor awareness.

## Conclusion

The overall baseline knowledge levels were very low and mean level of awareness on various broad issues (categories) of cervical cancer was 8.77. After education intervention knowledge levels improved to 30.39. Baseline knowledge on risk factors, screening and symptoms were better over HPV and vaccination. However, after the intervention, knowledge levels improved in all domains with low improvement about vaccination. Only 10 (3.5%) participants’ family members were ever screened for cervical cancer, although the 46.3% of them were willing to undergo screening, the important obstacle cost. Majority of the respondents did not hear the availability of vaccine and its primary preventive role in improving the risk of HPV infections. The result of this study revealed that only 33.9% of women had sufficient knowledge of cervical cancer after education intervention. This study also showed that a small percentage of study participants (9.9%) had an unfavorable attitude to seek medical help when they may any symptom of cervical cancer. The study also revealed that branch of study, year of study were significantly associated with knowledge levels of the students. Based on the findings of this study, education intervention is an effective method to improve knowledge levels on cervical cancer and students can be trained to disseminate the knowledge in society and help in spreading the positive attitude towards screening and vaccination. Based on this, we recommend the government should take measures to initiate health education training on cervical cancer at university levels and make educational institutions become important stakeholders to disseminate cervical cancer awareness and positive attitude in society.

## Acknowledgment

We thank all women students of Twedrows and Maraki campuses of the University of Gondar who took part in the survey. We also thank Institute of Biotechnology, University of Gondar, Gondar, Ethiopia for invaluable suggestions and support.

## Author Contributions

Conceptualization: NB, IM. Data collection tool (English): NB, IM & Data collection tool (Amharic): TM. Performed the experiments: IM, TM. Resources and supervision: NB. Analyzed the data: IM, NB, TM. Wrote original draft: IM. Editing & finalizing the original draft: NB, TM & IM.

## Supporting information captions The table in the Appendix

**Table A1:** McNemar test of various knowledge levels grouped as no, poor, fair, and good cervical cancer awareness about symptoms, risk factors, HPV, screening and vaccination of respondents at P=0.000 significance level.

| Domain | N=283/<br>McNemar<br>Value |
| --- | --- |
| Knowledge levels compared |  |
| Symptoms | 215.18 |
| Risk factors | 163.63 |
| HPV | 199.68 |
| Screening | 202.88 |
| Vaccination | 190.52 |

**Table A2:**
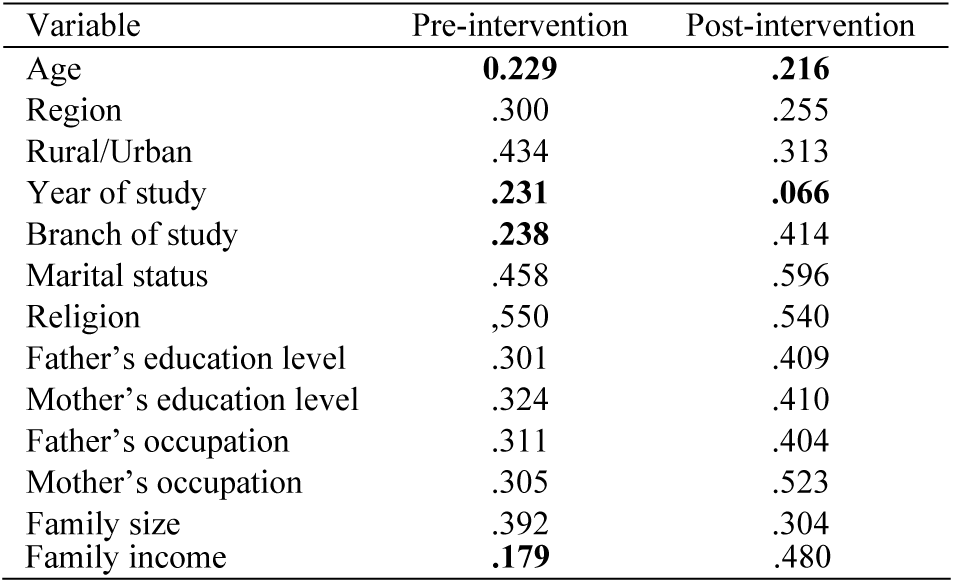
Significance of effect of various individual explanatory variables on overall awareness levels about CC.

**Table A3:** Various predictors relationship about CC at the P=0.05 significance level.

| Predictors relationship | Chi-square | Cramer's V |
| --- | --- | --- |
| Age category and Year of study | 103.82 | .606 |
| Age category and Father's education level | 14.82 | .229 |
| Age category and Mother's education level | 7.04 | .158 |
| Year of study and Branch of study | 26.01 | .303 |
| Branch of study and Family size | 7.47 | .163 |
| Father's education level and Mother's education level | 230.87 | .639 |
| Father's education level and Family size | 18.57 | .256 |
| Mother's education and Family size | 23.27 | .287 |

